# Development of a monkey avatar to study social perception in macaques

**DOI:** 10.1101/758458

**Authors:** Vanessa A.D. Wilson, Carolin Kade, Sebastian Moeller, Stefan Treue, Igor Kagan, Julia Fischer

## Abstract

Following the expanding use and applications of virtual reality in every-day life, dynamic virtual stimuli are of increasing interest in cognitive studies. They allow for control of features such as gaze, expression and movement, which may help to overcome limitations of using either static or poorly controlled real stimuli. In using virtual stimuli however, one must be careful to avoid the uncanny valley effect - where realistic stimuli can be perceived as eerie, and induce an aversion response. At the same time, it is important to establish whether responses to virtual stimuli mirror responses to depictions of a real conspecific. In the current study, we describe the development of a new avatar with realistic features for nonhuman primates, the ‘primatar’. As a first step towards validation, we assessed how monkeys respond to images of this avatar compared to images of real monkeys, and an unrealistic avatar. We also compared responses between original images and scrambled as well as obfuscated versions of these images. We measured looking time to images in six free moving long-tailed macaques (*Macaca fascicularis*) and eye movement exploration behaviour in three rhesus macaques (*Macaca mulatta*). Both groups showed more of such signs of overt attention to original images than scrambled or obfuscated images. In addition, we assessed whether the realistic avatar created an uncanny valley effect through decreased looking time, finding that in both groups, monkeys did not differentiate between real, realistic or unrealistic images. These results provide support for further development of our avatar for use in social cognition studies, and more generally for cognitive research with virtual stimuli in nonhuman primates. Future research needs to shed light on the source of the inconsistent findings for the uncanny valley effect in macaques, to elucidate the roots of this mechanism in humans.

## Introduction

The use of virtual reality is on the rise, and has been applied across a broad range of settings, from education and training to tourism and health (1–3). Recently, the development of virtual stimuli has been applied to social cognition research, by providing a life-like, social stimulus that can move in a controlled manner (4,5). The use of images and video footage of social content has been common place in social cognition studies, yet assessment of social responses with these stimuli can be limited. For example, static stimuli lack movement which can quickly reduce the realism in their appearance (6). Whilst video footage can counter this limitation (7), collecting footage that fits exact experimental requirements can be challenging. As cognitive studies seek to answer more detailed questions about perception and response to social stimuli, the introduction of virtual stimuli to cognitive research could help to resolve current methodological limitations in understanding social cognition and perception.

What is particularly valuable about this approach is the ability to control exactly the gaze, expression and movement of the stimulus, something which is limited when presenting footage of real animals. It is important, however, in developing stimuli of a realistic nature, to avoid the uncanny valley effect. In humans, as a stimulus becomes more realistic in its appearance, human affinity also increases, up to a point where a particularly ‘realistic’ stimulus creates an aversion response – known as the uncanny valley (Figure 1).

**Figure 1.**
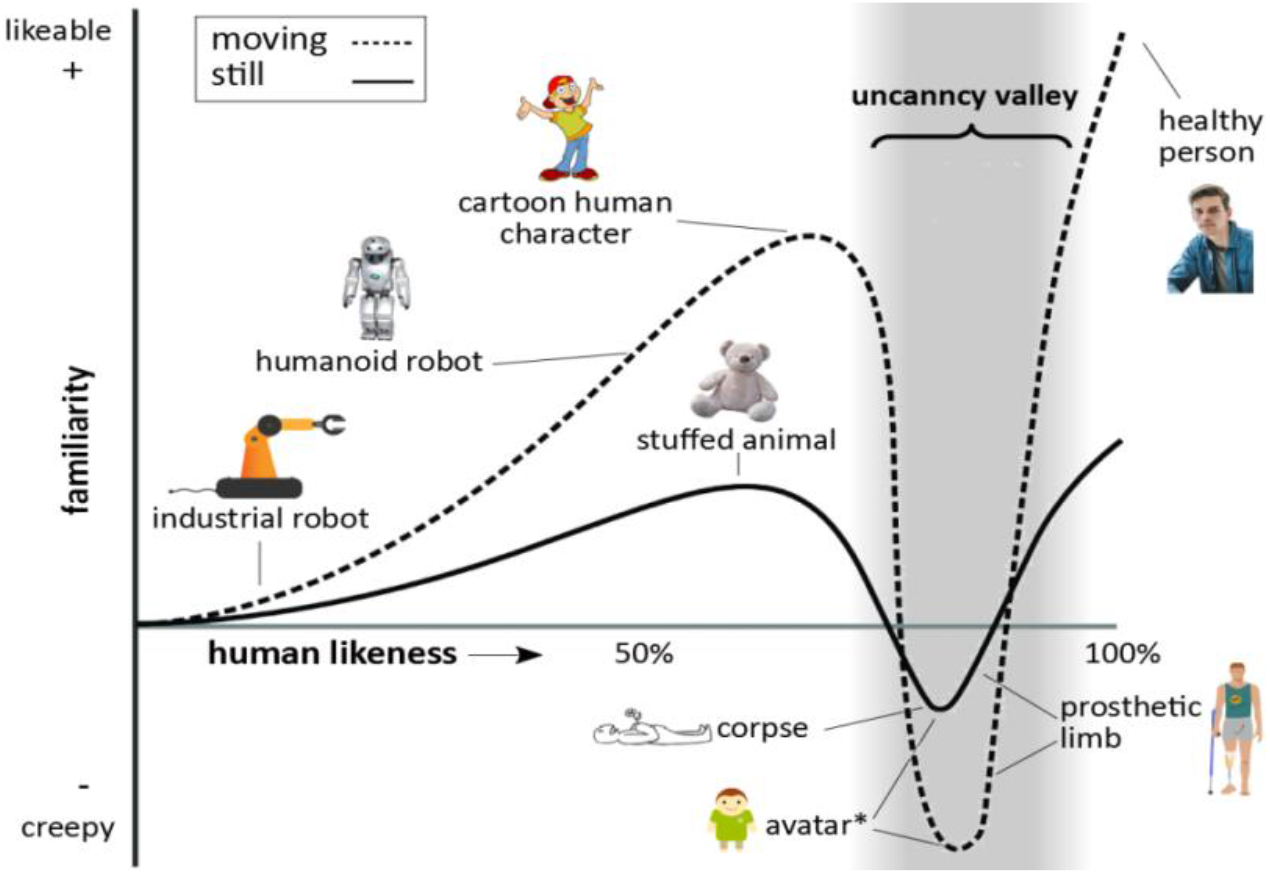
The uncanny valley. Figure reflects features as described by (8,9,17). The trough denoted by asterisk relates to findings by Steckenfinger & Ghazanfar (17), who found an uncanny valley effect in long-tailed macaques for both static and dynamic virtual stimuli.

Since the uncanny valley theory was first posited by Mori (8,9), there have been numerous attempts to test this theory (4,10) and to understand what mechanisms might underlie this response (11,12) including examining the neural mechanisms (13). There are currently two main competing theories. One theory is that aversion to ‘eerie’ stimuli is acquired, occurring through development. For example, eeriness ratings of a super-computer increase with perceptions of the machine’s ability to experience human emotions (14). Children under nine years old did not perceive a human-like robot as creepier than a machine-like robot, while children over nine years old found the human-like robot creepier, indicative of an uncanny valley effect emerging with age (15).

The alternative explanation is that the uncanny valley is an innate response, shaped by survival of people who were more likely to avoid potentially dangerous stimuli. For example, avoidance of aversive facial aesthetics (4,10), disgusting stimuli (16) or corpses (11,12) could provide a survival advantage, providing future generations with aversive responses to stimuli that elicit feelings of danger. Evidence for the uncanny valley in long-tailed macaques (*Macaca fascicularis*) supports the notion that this is an evolved response. Monkeys (N=5) showed decreased viewing time to a ‘realistic’ avatar over an unrealistic avatar or real monkey stimulus (17), suggesting that aversive responses to potentially dangerous stimuli may have occurred in a common primate ancestor. Whilst this evidence is limited, it does suggest that researchers should take care when creating virtual stimuli for nonhuman animals, in case they inadvertently create a stimulus that study subjects perceive as eerie.

Several studies have recently assessed social responses in nonhuman primates using virtual faces (5,18–20), yet the influence of realism of these stimuli on evoking visual exploration responses has not been addressed. In the current paper, we report the development of a new more realistic avatar stimulus, for use in social cognition experiments. Here, we 1) describe its development and 2) compare gaze responses of both rhesus (*Macaca mulatta*) and long-tailed macaques to a static version of the avatar, in relation to real, unrealistic and scrambled stimuli, thereby testing for avoidance effects of the realistic stimulus. We predicted that monkeys would look longer at the facial images than the scrambled images. We also predicted that if the realistic avatar was perceived as eerie, we would see a decrease in looking time towards the realistic avatar compared with the real images or unrealistic avatar. We additionally tested response to obfuscated versions of the different stimuli. We predicted that, as the obfuscated images reduce image detail, looking times to images would decrease with stronger obfuscation.

## Methods

### Avatar development

We developed an avatar based on measurements and features of a female long-tailed macaque. The graphical features of the avatar were developed in Blender (version 2.79b) using Archlinux OS, with six main phases. 1) 3D modelling, which determined the shape and dimensions of the skull. To establish head proportions, we used images and 3D scans of adult female long-tailed macaque skulls, and followed the facial proportions provided by Schillaci et al. (21) 2) 3D sculpture of skin details, which allows one to add details such as skin tint and wrinkles. To create realistic details we examined photographs and footage of female long-tailed macaques housed at the German Primate Centre. 3) Skin and eye texturing. This creates ‘realistic’ features of eyes and skin by adding in facial shading and details such as eye colouration and reflections. 4) Development of facial rig which provides the underlying structure for movement of eyes, eyebrows, nose, jaw and cheeks. 5) Facial animation, which allows facial expressions to be formed from adjustments of the facial rig. 6) Addition of hair. Hair textures were created in Blender, with varying shades and lengths. To add hair to the 3D model, the mesh of the skin is unwrapped into a 2D surface, allowing the skin texture to be painted with the hair. This is necessary to ensure that the hair texture is fully aligned with the skin vertices. For the current study, we tested a static prototype with neutral expression after phase 3, prior to further development (Figure 2).

**Figure 2.**
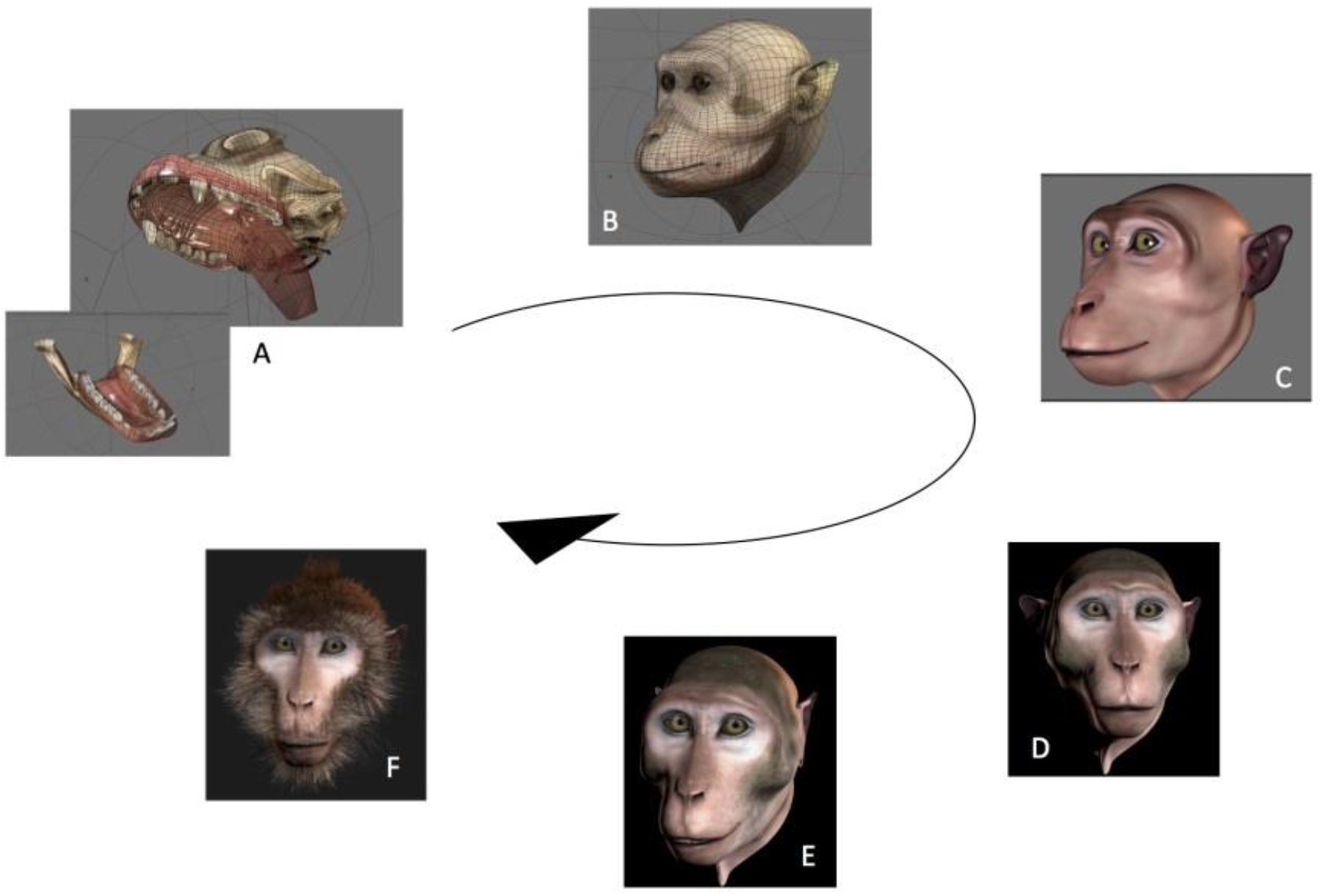
Development of the avatar. Letters denote modelling of the skull dimensions (A), sculpture and texturing of the skin (B-D), development of facial rig and animation (E) and addition of hair (F). For the current experiments we used the avatar from stage D.

#### Eye tracking study

*Subjects*. Three male rhesus macaques (*Macaca mulatta*) between 8 and 10 years old were tested. Monkeys were pair-housed with other males at the German Primate Center, Göttingen, in two enclosures (23.3m^3^ and 21.3m^3^) conjoined by an overhead tunnel. Individuals participated daily in testing in a separate room and were rewarded for participation with water and juice. All animals had been previously implanted with cranial plastic “headposts” under general anaesthesia and aseptic conditions, for participating in neurophysiological experiments. The surgical procedures and purpose of these implants were described previously in detail (22). Monkeys were previously trained in both using the primate chair, and in fixating their gaze for eye tracking calibration.

##### Stimuli

Stimuli consisted of three images of real long-tailed macaques (adult females), one image of a realistic avatar and one image of an unrealistic avatar (Figure 3), each presented on a black background. All images had averted gaze. Images were scaled in GIMP to 673 × 780 pixels (~22 × 25 degree visual angle). For each image, a block-scrambled (17 × 20 pixel), mildly obfuscated (35% pixelation, 25 pixel blurring radius) and strongly obfuscated (70% pixelation, 50 pixel blurring radius) version was also presented (Figure 3). Image scramble and obfuscation was conducted directly within the experimental software.

**Figure 3.**
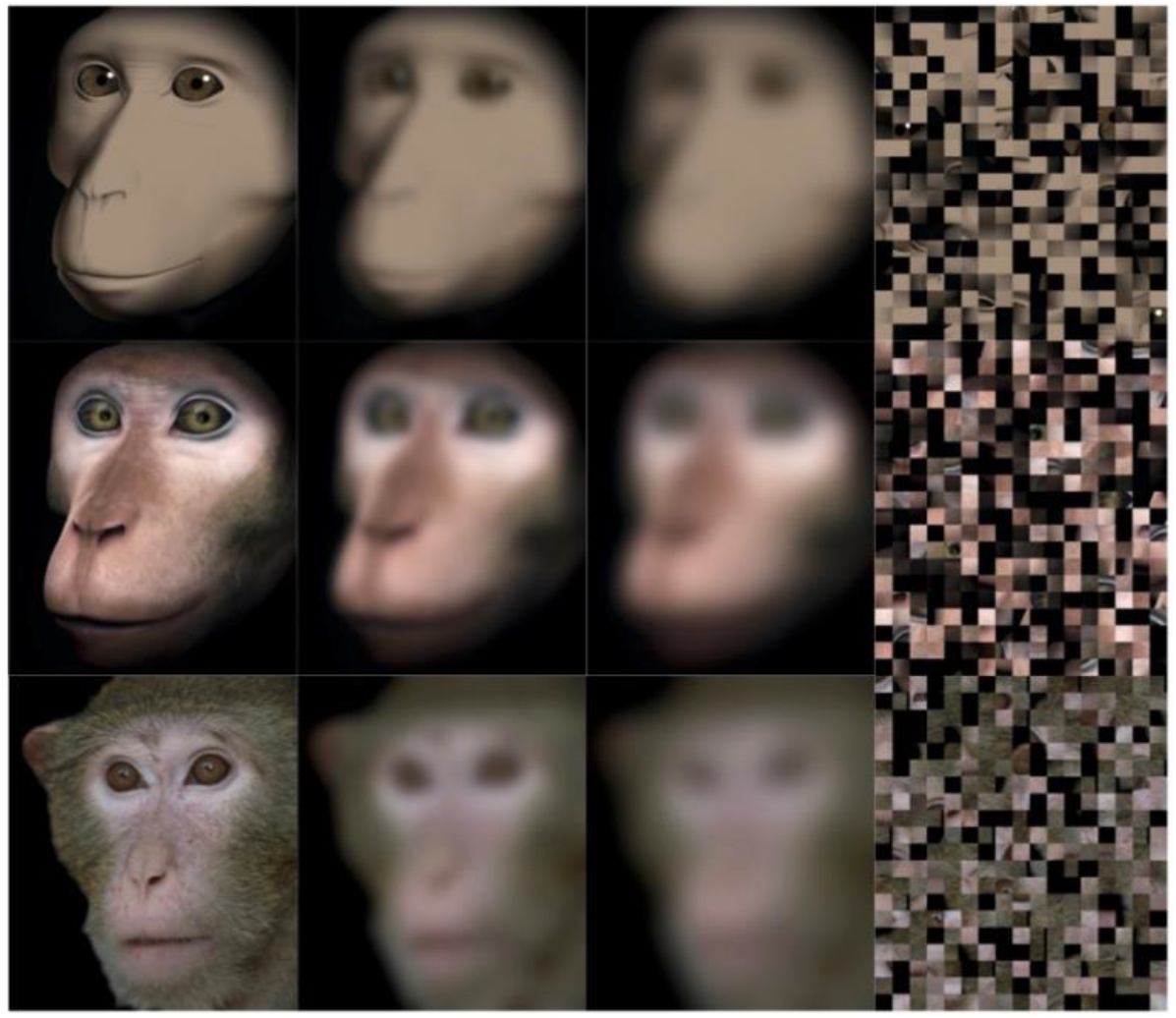
Test stimuli. Top to bottom: unrealistic avatar, realistic avatar, and real monkey (one of three real photographs used). Left to right in each row: original image, mild obfuscation, strong obfuscation, scrambled image.

##### Apparatus and procedure

Monkeys were head-fixed in a primate chair placed in front of a visual display (Set-up 1: 55” diagonal, EYE-TOLED-5500, Eyevis, Reutlingen, Germany; Set-up 2: 27” screen Acer HN274H, Acer Computer GmbH, Ahrensburg, Germany). Experiments were run using EventIDE (Okazolab, Delft the Netherlands). Eyetracking was performed using camera-based infra-red tracker (Set-up 1: EyeLink 1000+, SR-Research, Ottawa, Ontario, Canada; Set-up2: MCU02 ViewPoint, Arrington Research, Scottsdale, Arizona, USA). Each session consisted of 75 trials. Each trial started with a red dot inside a yellow circle (for 1500 milliseconds), on which the monkeys had to fixate to receive a drop of liquid. They were then presented with a scrambled image for five seconds. This was followed by a second fixation, and a non-scrambled image for five seconds. Each real monkey image was presented five times each, and each avatar image was presented fifteen times each, giving a total of 15 presentations per condition. Each obfuscated image was presented three times each. One monkey participated in one session, and two monkeys in two sessions. We measured total fixation time per facial image.

##### Ethics

The experimental procedures were approved of by the responsible regional government office (Niedersächsisches Landesamt fuer Verbraucherschutz und Lebensmittelsicherheit (LAVES)^1^).

##### Analyses

Eye-tracking data were extracted and analysed using MATLAB (version 2014b). When monkeys participated in two sessions, we analysed data from only the first session, as apparent attention to the images decreased during repeated sessions, with fewer fixations making data analysis unreliable. Individual gaze responses are shown in Figure 4. We plotted proportions of fixation durations per image, to each image type. We created average heat maps for each scrambled image and for each original image, and created bar plots of proportion fixation durations to the whole face image, eyes and mouth regions, for each image.

**Figure 4.**
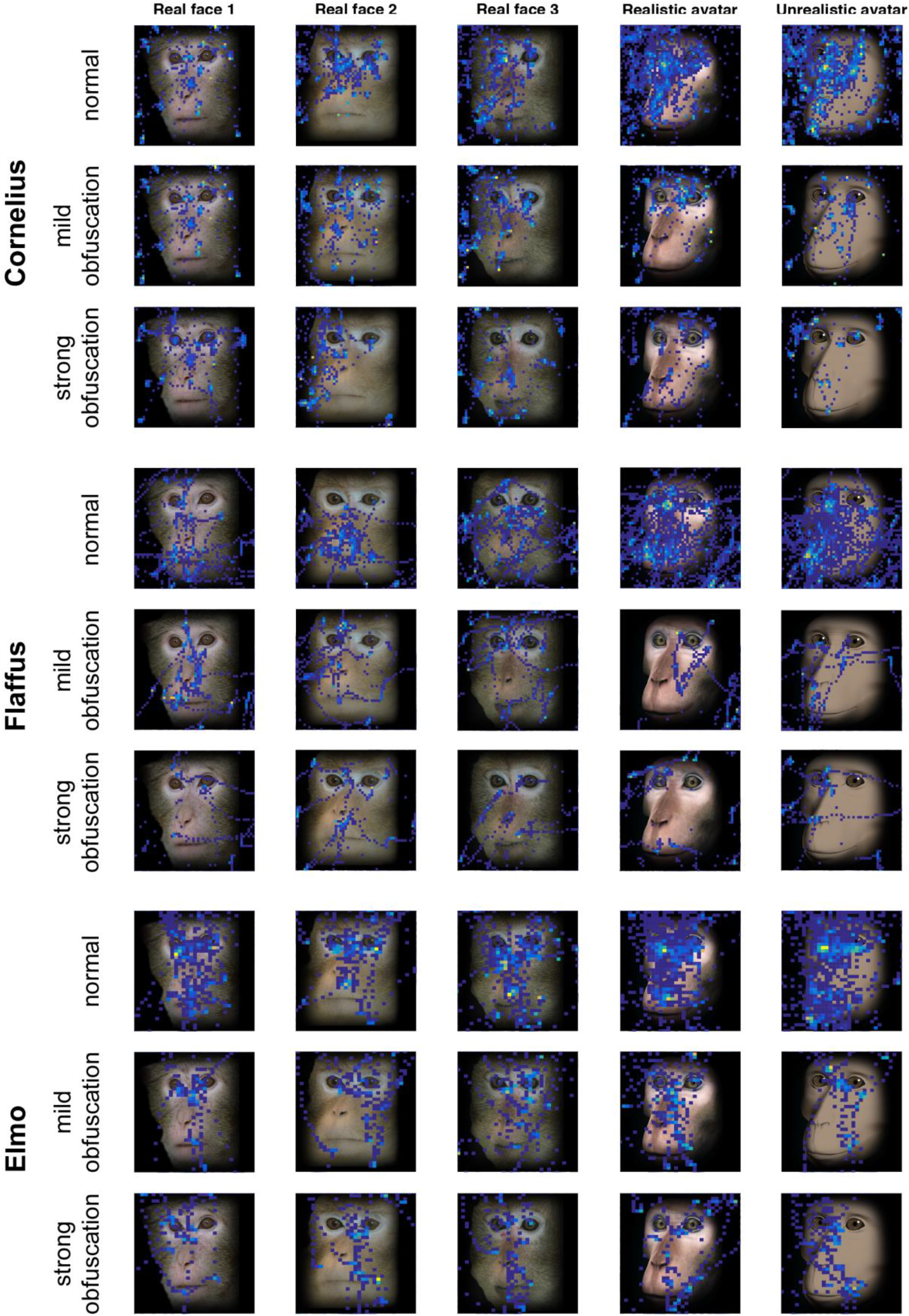
Total gaze fixations to real monkey and avatar images per monkey.

Statistical analyses were conducted in R Studio (version 1.0.153) using linear mixed models and running bootstrapped confidence intervals for each model. Whilst we wished to analyse data for each monkey separately, this led to problems with model convergence, thus we conducted our analyses across all three subjects, treating ID as a random variable in all models. All dependent variables were scaled as Z-scores prior to analysis. Alpha was set to .013 to correct for multiple comparisons between four different models.

In our models, we first examined whether the monkeys exhibited similar patterns of attention to images across different obfuscations (prediction 3). We examined total gaze fixation time to the image as the dependent variable, with obfuscation type as the fixed variable. We compared looking time to original (unobfuscated) images and strongly obfuscated images against mildly obfuscated images.

To test the first prediction, we examined whether attention (as measured by duration of eye fixation) to the images differed between the scrambled compared with original images, across both real monkey photographs and avatar images.

To test the second prediction, we examined fixation time to each image between the original real monkey and avatar conditions: specifically, we compared the unrealistic avatar and real monkey images against the realistic avatar. To correct for multiple comparisons between models, for total fixation time alpha was set to 0.017 (3 models), and for face fixation time alpha was set to .013 (4 models).

#### Looking time study

##### Subjects

Ten long-tailed macaques were trained on the task, and six reached criterion and were tested (3F, 3M, 2 – 10 years). One female only completed half of the test sessions (block 1). Participants belonged to a group of 36 captive-housed long-tailed macaques (*Macaca fascicularis*), housed at the German Primate Center, Göttingen. The group had access to both an indoor (49 m^2^) and outdoor (141 m^2^) enclosure, with ad libitum access to food and water as well as enrichment. Individuals participated voluntarily in all cognitive testing, which took place in a separated indoor testing area which could be divided into six cubicles (2.6 m × 2.25 m × 1.25 m; h × w × d). Testing hours were from 10:00 – 12:00 and 14:00 – 18:00, Monday to Friday. Monkeys were rewarded for participation during testing with cut raisins.

##### Stimuli

To train the monkeys on a viewing paradigm, we selected sixty novel images of non-social content. Images were divided into three categories - landscape, object and food, content which was deemed motivating for the monkeys to view. Images were scaled in GIMP to dimensions of 4000 × 3000 pixels. Test images were the same as for the eye tracking study (Figure 3). However, to reduce the number of trials, we presented only two images of real monkeys. Block-scrambled images were formed prior to testing using Matlab (version 2018b). Images were obfuscated in GIMP using the pixilation and pixel Gaussian blur infinite impulse response filters (mildly obfuscated = 5 pixel pixilation and 45 pixel Gaussian blur; strongly obfuscated = 10 pixel pixilation and 90 pixel Gaussian blur).

##### Apparatus and procedure

The monkeys were tested on an Elo 17” SXGA TFT touchmonitor which was connected to an external MacBook Pro computer which ran on OS X El Capitan version 10.11.6. In this set up, cameras from the side and above filmed the monkeys. Experiments were run using MWorks (version 0.7, https://mworks.github.io). Each trial consisted of an image presented on screen (19.63cm × 19.63 cm) with a touch-target beneath the image (white square: 3.93 cm × 3.93 cm). The monkeys could view the image for 60 s, or, by touching the target, change the image sooner. Each training session consisted of 20 trials. Reward was given at random intervals, so monkeys were not reinforced to touch the target, thereby removing food-based incentives for viewing the images. Order of image presentation was randomized. To reach criterion, monkeys had to touch the target on each trial over one session.

For the testing procedure, monkeys received two blocks of stimuli. This design was to reduce the attentional demands on the subjects, who easily lost motivation, by presenting fewer trials per test session. Block 1 consisted of original images (real monkey, realistic avatar, unrealistic avatar) and scrambled images. Block 2 consisted of obfuscated images only. Block 1 was presented to all subjects first, as in case the monkeys lost interest, we wished to prioritise the original over the obfuscated images. Each block consisted of three sessions, and each session was 8 trials long. Between each session, monkeys were given a new training session to reduce their expectation of social stimuli in the next test session (except in one case: Linus, missing one in-between training session). We measured looking time per image by coding looking time from videos of each test session. Monkeys were considered to be attending to the images when their head and eyes were clearly oriented towards the screen.

##### Ethics

This study was non-invasive and is in accordance with the German legal and ethical requirements of appropriate animal procedures using nonhuman primates. As confirmed by the competent authority (the Niedersächsisches Landesamt für Verbraucherschutz und Lebensmittelsicherheit (LAVES)), these experiments do not constitute a procedure according to the animal welfare legislation (§7, Abs. 2 TierSchG); therefore a permit was not required (LAVES document 33.19-42502-04). Institutional approval was provided by the German Primate Center Animal Welfare Body (application no. E2-18).

##### Reliability coding

All test videos were blind coded by a coder naïve to the hypotheses of the study. For reliability assessment, ten videos were coded by VW who was not blind to the hypotheses. Due to a missing image change in one file, reliability was calculated for nine videos using Spearman’s *rho*. Looking time showed good reliability between two coders, for nine videos (*rho* = 0.84).

##### Analyses

Analyses were conducted in R Studio (version 1.0.153). We ran mixed effects generalized linear models, with a Gamma distribution, using the lme4 package. We set the number of adaptive Gauss-Hermite quadrature points to zero to aid model convergence. For all monkeys, looking time was examined as the dependent variable, image type as a fixed effect and ID as a random effect. As looking time was highly skewed, we analysed looking time that had been transformed using a Yeo Johnson transformation. Trials where monkeys were judged not to look at the images were removed from analysis. For all models, we ran bootstrapped 95% confidence intervals. Alpha was set to .013 to correct for multiple comparisons between four different models.

In model 1, we compared looking time to the scrambled images with looking time to images of original real monkeys and avatars combined. In model 2, we compared looking time between original realistic avatar and real monkey images, and between realistic avatar and unrealistic avatar images. In model 3, we compared looking time to obfuscated social images over non-obfuscated social images. We also wished to examine looking time between realistic avatar, unrealistic avatar and real monkey images within both mild and strong obfuscation categories; however, as the monkeys often did not look at these images, the sample size was too small to examine differences within obfuscation type. Instead, in our final model, we examined whether the monkeys’ gaze differentiated between the different levels of obfuscation, that is, by grouping looking time to avatar and real monkey images within mild, and within strong, obfuscation.

## Results

### Eye tracking study

Fixation time on the face images differed with obfuscation, i.e. monkeys looked longer at original (unobfuscated) as compared to mildly obfuscated images (*b* = 0.50, *SE* = 0.16, *p* < .01, 95% CI = [0.22, 0.82]) and looked longer at mildly obfuscated as compared to strongly obfuscated images (*b* = −0.41, *SE* = 0.19, *p* < .05, 95% CI = [−0.76, −0.06]), suggesting that the reduced image detail of obfuscated images weakens interest (Figure 4).

In keeping with our first prediction, the rhesus macaques looked less overall at scrambled as compared to non-scrambled images (*b* = −1.27, *SE* = 0.10, *p* = < .001, 95% CI = [−1.45, −1.10]), and looked more at the face regions of original images as compared to the corresponding regions in scrambled images (*b* = −0.42, *SE* = 0.10, *p* = < .001, 95% CI = [−0.67, −0.27]). Figure 5 indicates increased fixation around the eyes and mouth region of the original images compared with the scrambled images. Regarding our second prediction, we found no difference in time spent attending to the original faces between the unrealistic avatar and the realistic avatar (*b* = 0.06, *SE* = 0.19, *p* = 0.73, 95% CI = [−0.30, 0.42]) or between the real monkey and realistic avatar images (*b* = −0.19, *SE* = 0.18, *p* = 0.28, 95% CI = [−0.56, 0.13]). The monkeys therefore did not appear to direct their gaze to the realistic avatar any less than they did to the other images, providing no evidence that the realistic avatar created an uncanny valley effect (Figures 5,6).

**Figure 5.**
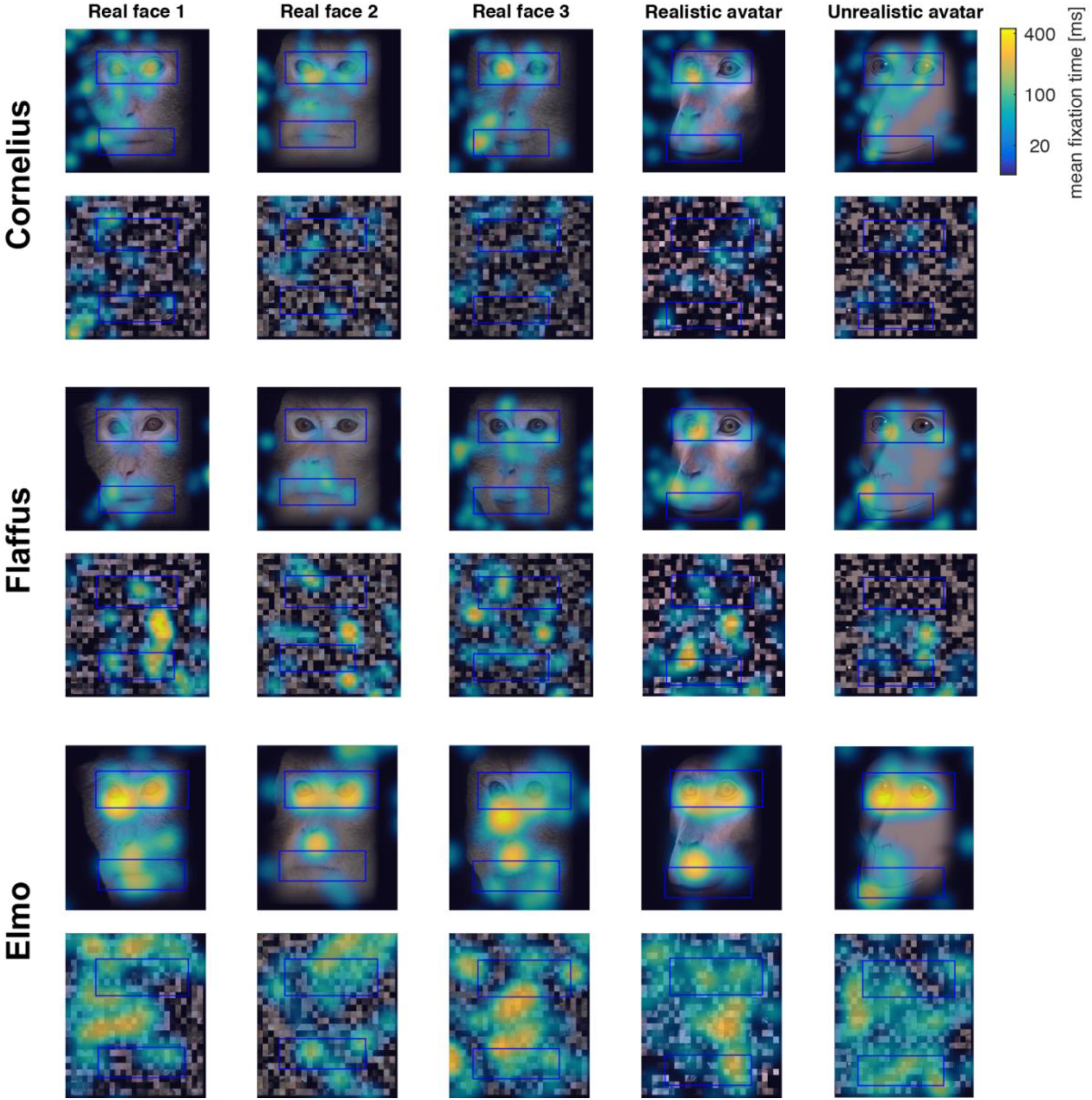
Average gaze fixation time to original and scrambled images across three monkeys.

**Figure 6.**
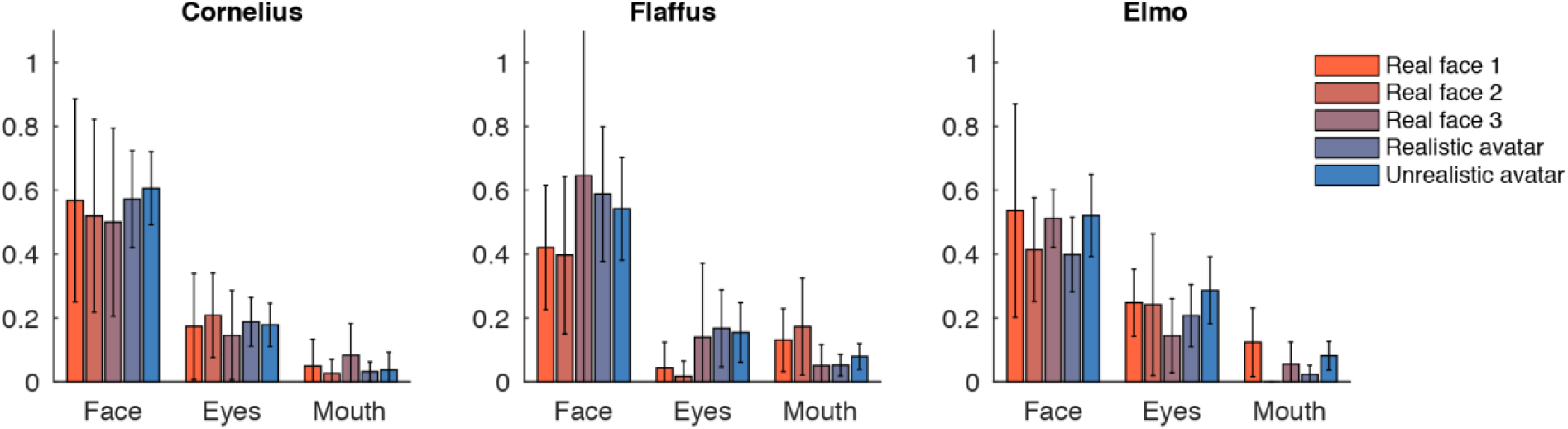
Gaze fixations of three monkeys to the face, eyes and mouth of original real monkey and avatar images. Eye and mouth regions are defined by the blue boxes visible in Figure 5. Error bars indicate confidence intervals.

### Looking time study

Similar to findings for the rhesus macaques, the long-tailed macaques looked at original facial images significantly more than scrambled images, as predicted, although the confidence intervals suggest this difference to be only marginal (*b* = −0.59, *SE* = 0.13, *p* < .001, 95% CI = [−1.26, 0.12]; see Figure 7). There was no difference in looking time between the realistic avatar images, and either the real monkey (*b* = 0.10, *SE* = 0.20, *p* = 0.61, 95% CI = [−1.00, 1.24]), or the unrealistic avatar (*b* = 0.13, *SE* = 0.23, *p* = 0.57, 95% CI = [−1.39, 1.43]; see figure 7). This again provides no evidence for an uncanny valley effect of the realistic avatar. For our third prediction, we found that the monkeys did look less overall at obfuscated social images than original social images (*b* = −0.75, *SE* = 0.12, *p* = < .001, 95% CI = [−1.48, −0.01]), although they did not differentiate between mild and strong obfuscations of social images (*b* = −0.04, *SE* = 0.17, *p* =0.82, 95% CI = [−1.02, 0.83]).

**Figure 7.**
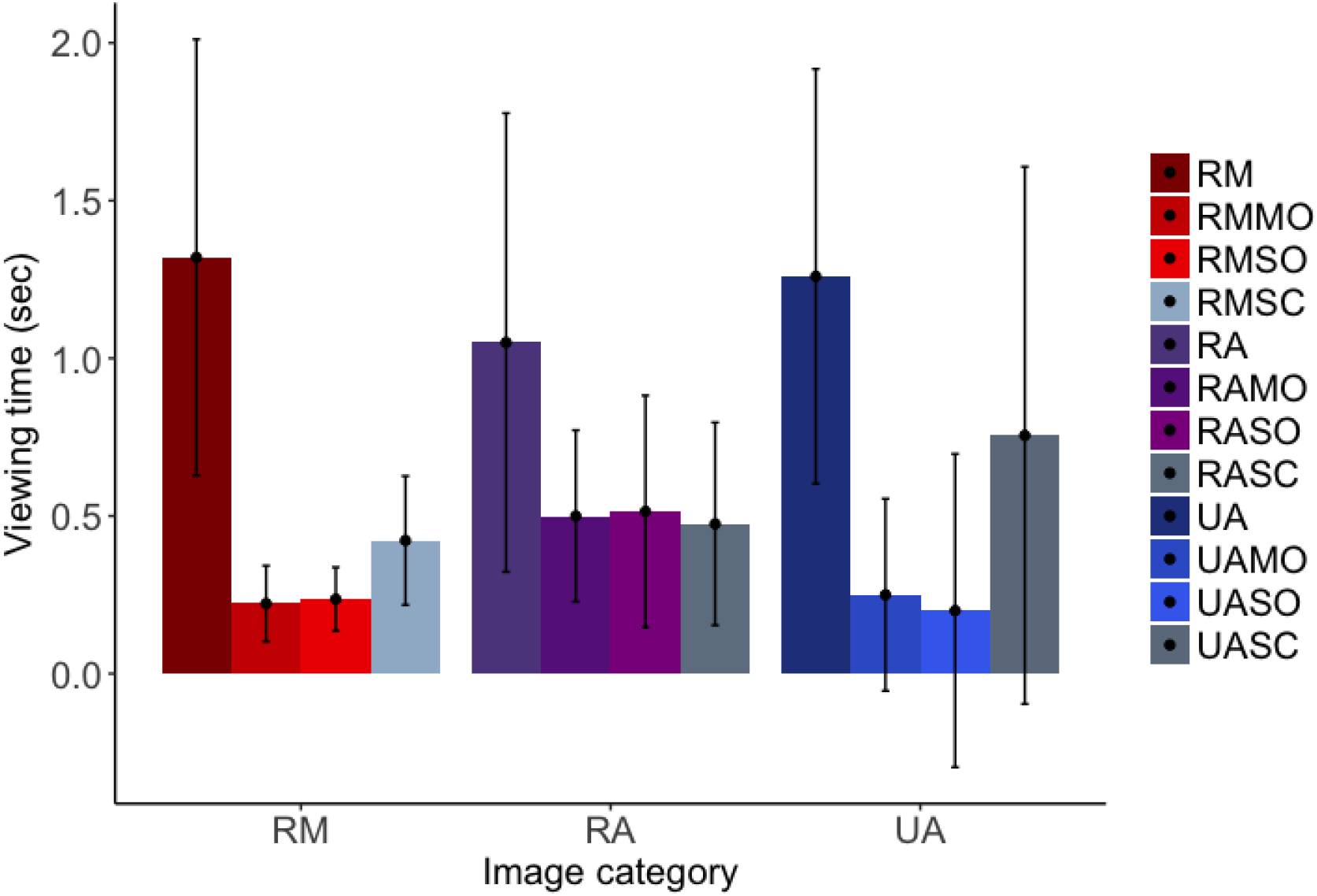
Long-tailed macaques’ looking time to each image category. RM = real monkey, RA = realistic avatar, UA = unrealistic avatar, MO = mild obfuscation, SO = strong obfuscation, SC = scrambled image. Error bars indicate 95% confidence intervals.

## Discussion

Our results, from both long-tailed and rhesus macaques, indicate that our realistic avatar does not cause an uncanny valley effect, i.e. an aversion of overt attention. This finding raises two possible interpretations. Firstly, addressing hypotheses for the uncanny valley, our results, in contrast to Steckenfinger & Ghazanfar (17), do not support the theory that this aversive response to certain social stimuli occurred in a common primate ancestor. Whilst Steckenfinger & Ghazanfar (17) assessed responses to not only static but also dynamic stimuli, one limitation of their study was that the realistic avatar was in colour, whilst the unrealistic avatar was black and white. This could explain why the monkeys’ attention to the two different stimuli varied. An alternative interpretation is that in the current study, the avatar we created was realistic enough not to create an aversion effect, whereas in the prior study, the realistic stimulus contained some aversive features which reduced subjects’ attention. To fully establish whether nonhuman primates do exhibit an uncanny valley effect, further research is necessary that examines responses to stimuli along a continuous gradient from unrealistic to realistic. Doing so would also clarify at what point certain features become aversive.

Concerning the monkeys’ aversion and attention to virtual stimuli, our results, which indicated no difference in gaze allocation between real and realistic images, support the use of our virtual stimulus to assess social interactions and behaviours in macaques. The eye tracking results, in particular, indicate similar patterns of attention to both the real and avatar faces, reflecting previous findings that monkeys attend primarily to the eyes, followed by the nose and mouth regions (23–25). As the avatar that we used here was a prototype for further experiments, our current findings support further development of the avatar, as described in figure 2. These results also suggest that, at least for static images, differences in facial features such as skin texture and eye colour might not be so important for virtual stimuli. We suggest that further investigation of the role of ‘realistic’ features in attention-aversion requires further investigation, especially for dynamic stimuli.

Measures of attention continue to be crucial to studies of social cognition. Recent studies however, have raised concerns about the presentation and type of stimuli used in these studies (6,7), as well as the interpretations of attention bias (26,27). Virtual stimuli may therefore provide an alternative to traditional static stimuli, countering issues such as lack of movement or facial expression, as well as providing a method to better interpret social attention. For example, a virtual stimulus may allow for manipulations of differences in facial features such as gaze direction (28), emotional expression (29,30), sex (31,32), age (33), symmetry (34,35), status (31,36,37) and assertiveness (38), features which provide important social information and therefore may influence conspecific attention. Murphy & Leopold (5) recently demonstrated face-selective neurons in rhesus macaques respond to static images of a virtual monkey, and that neural responses varied with certain variables, such as head orientation and emotional expression. The use of avatar stimuli could allow for greater control over subtle feature differences found in non-virtual stimuli such as photographs, and thereby reduce noise in the data, allowing for better interpretation of social preferences and naturalistic responses.

Our study is not without limitations. Specifically, the decreased attention shown by the long-tailed macaques to the obfuscated images could be accounted for by order effects, as the obfuscated images were viewed in separate sessions after the original (unobfuscated) images. A decrease in image novelty could account for this reduced attention, however, results from the rhesus macaques do indicate that, regardless of order, attention to social images decreases with obfuscation. An additional limitation is our small sample size in both species. Despite this, our results, which come from two different species in different testing environments and using slightly different test procedures, converge on the finding that the realistic avatar does not create an uncanny valley effect. This is an important finding worthy of further investigation in nonhuman primates.

## Acknowledgements

The authors acknowledge funding from the Ministry for Science and Education of Lower Saxony and the Volkswagen Foundation through the program “Förderung exzellenter Forschung in Niedersachsen: Forschungsverbund” Zuwendungsnummer VWZN3106, as well as the Leibniz Association through funding for the Leibniz ScienceCampus Primate Cognition and support for the project provided by a seed fund. The authors would also like to acknowledge Damien Monteillard for graphical design, and Ilia Korjoukov of Okazalab for providing programming support. For providing long-tailed macaque skulls for 3D modelling we acknowledge Kerstin Maetz-Rensing, and for 3D scans of long-tailed macaque skulls we thank Borna Mahmoudian. We are indebted to Anton Unakafov for developing the MATLAB script and to Heidrun Otto for video coding. We also thank Benjamin Walter for the art work and for providing technical advice. Finally, thanks to Ralf Brockhausen for technical assistance, and the animal caretakers, Nadia Rabah and Henning Mascher, for their support.

## Authors’ contributions

VW, SM, IK, JF and ST conceptualized and designed the eyetracking task. SM and IK implemented the eyetracking task and conducted the experiments. VW and SM analysed the eyetracking data. VW, CK and JF designed the looking time study. VW and CK conducted the looking time study. VW and SM prepared the figures. VW wrote the initial version of the manuscript. All authors discussed and interpreted the findings, and revised the manuscript.

## Footnotes

1 Lower Saxony State Office for Consumer Protection and Food Safety

